# Novel selective CDK4 inhibitor ALK-13743: Functional determinants of response in ER+ breast cancer models

**DOI:** 10.64898/2026.11.17.767343

**Authors:** Vishnu Kumarasamy, John Sherrill, Ryan Henry, April Bruno, Jordan Bartlebaugh, Jianxin Wang, Thomas N. O’Connor, Agnieszka K. Witkiewicz, Amy Hart, Andy Phillips, Kiel Lazarski, Eunice Park, Erik S. Knudsen

**Affiliations:** Department of Molecular & Cellular Biology, Roswell Park Comprehensive Cancer Center, Buffalo, NY, USA 14263; Aleksia Therapeutics, 11757 W. Ken Caryl Avenue, Suite F, 520, Littleton CO, USA 80127

**Keywords:** CDK4, CDK4/6 inhibitor, breast cancer, cell cycle, RB1, PI3K, ALK-13743, atirmociclib, palbociclib

## Abstract

Cyclin-dependent kinase 4 (CDK4) and CDK6 are central drivers of cell cycle progression in estrogen receptor-positive (ER+) breast cancer, and pharmacological inhibition of this pair has become a standard-of-care therapeutic strategy. However, currently approved CDK4/6 inhibitors—palbociclib, ribociclib, and abemaciclib—carry associated toxicities, including myelosuppression and other adverse effects which are associated with inhibition of CDK6 that limits tolerability and combinatorial strategies. Here we describe the preclinical characterization of ALK-13743, a novel, highly selective CDK4 inhibitor. ALK-13743 potently inhibited CDK4/Cyclin D1 with an IC50 of 33 nM and displayed selectivity over CDK6/Cyclin D3 (IC50, 99 nM), CDK2/Cyclin E1 (IC50, 1854 nM), and CDK1/Cyclin B1 (IC50, >10000 nM) in biochemical kinase assays. In cellular models, ALK-13743 demonstrated potent antiproliferative activity in CDK4-dependent ER+ breast cancer cells at nanomolar concentrations, while exhibiting limited inhibition in CDK6-dependent cell lines. ALK-13743 exhibited single-agent growth suppression and drug synergy with the selective estrogen receptor degrader giredestrant and PI3K pathway inhibitors but was entirely RB-dependent and blunted by CDK6 overexpression. Single-agent ALK-13743 treatment led to tumor regression in T47D xenografts *in vivo*. ALK-13743 is a potent, CDK4-selective inhibitor dependent upon RB for its antiproliferative profile in ER+ breast cancer models. Its favorable combinability with multiple clinically relevant targeted agents supports further development, particularly in combination regimens, and positions it as a differentiated CDK4-selective therapy for ER+ breast cancer.

## Introduction

Hormone receptor-positive (HR+), HER2-negative (HER2-) disease accounts for approximately 70% of all cases of breast cancer[1, 2]. The CDK4/6–RB axis is a critical regulator of the G1/S cell cycle transition in HR+/HER2-breast cancer[3–5]. Estrogen signaling functions as the primary mitogen for HR+/HER2-breast cancer and is the target of multiple endocrine therapies including aromatase inhibitors and selective estrogen receptor degraders. Estrogen signaling drives cell cycle entry in part by driving Cyclin D1 expression, leading to CDK4/6 activation and subsequent phosphorylation and inactivation of RB, enabling transcription of E2F target genes required for S-phase entry[6]. Numerous preclinical studies showed that HR+/HER2-breast cancers were particularly sensitive to catalytic CDK4/6 inhibitors, which supported clinical trials in this disease setting[7–9].

The use of CDK4/6 inhibition in HR+/HER2-breast cancer has been one of the most significant advances in breast oncology of the past decade[5, 10, 11]. Three CDK4/6 inhibitors— palbociclib, ribociclib, and abemaciclib—are approved in combination with endocrine therapy for metastatic breast cancer. These agents roughly double progression-free survival when combined with endocrine therapy, as compared to endocrine therapy alone. Additionally, abemaciclib and ribociclib are approved in the context of adjuvant therapy for high-risk HR+/HER2-breast cancer where inclusion of CDK4/6 inhibitors has shown to improve invasive disease-free recurrence[12, 13]. Despite this success, these therapies are not curative and all three approved agents harbor hematological toxicity, primarily causing neutropenia[4, 14–17]. This feature of CDK4/6 inhibitors is linked to CDK6 inhibition, as this kinase is particularly important in hematopoiesis[10, 18]. Recently published studies have illustrated that HR+/HER2-breast cancer models are particularly dependent on CDK4, but not CDK6, providing an important rationale for developing CDK4 - selective inhibitors[19]. Thus, selective CDK4 inhibition might decouple myelosuppression from antitumor efficacy, potentially improving the therapeutic window.

The investigational CDK4-selective agent atirmociclib (formerly PF-07220060) has provided preclinical support for this hypothesis[19]. Furthermore, both activity in HR+/HER2-breast cancer and an improved tolerability profile compared to dual CDK4/6 inhibitors has been presented[20, 21]. Here we describe the preclinical characterization of ALK-13743, a novel CDK4-selective small-molecule inhibitor developed by Aleksia Therapeutics. We demonstrate that ALK-13743 is a potent, selective CDK4 inhibitor that induces RB-dependent growth arrest and antiproliferative activity in ER+ breast cancer cell lines, confers broad combination activity with endocrine and PI3K-pathway-directed therapies in RB-proficient cells, and loses activity in settings of *RB1* loss or *CDK6* overexpression. These results support the development of drugs based off ALK-13743 for clinical use in RB-proficient HR+/HER2-breast cancer.

## Materials and methods

### Compound synthesis and characterization

ALK-13743 was synthesized by the Aleksia Therapeutics medicinal chemistry team. Compound identity and purity (>99%) were confirmed by HPLC-MS and NMR. Description of its synthesis can be found in its patent filing at https://patentscope.wipo.int/search/en/detail.jsf?docId=WO2025240517&_cid=P22-MWDYEA-20599-1[22].

### Biochemical kinase TR-FRET assays

A CDK time-resolved fluorescence resonance energy transfer (TR-FRET) assay was used to assess the potency of the compounds described herein. The assays were conducted using the following protein constructs: GST-CDK1 (M1-M297)/CCNB1 (M1-V433) (Carna Biosciences, Cat No 04-102), GST-CDK2 (M1-L298)/ CCNE1 (M1-A410) (Carna Biosciences, Cat No 04-165), GST-CDK4 (S4-E303)/ GST-CCND1 (Q4-I295) (ProQinase 0142-0143-1), GST-CDK6 (M1-A326)/6His-CycD3 (M1-L292) (ProQinase 0051-0373-1) and performed as follows.

Assays were performed in 384-well ProxiPlates (Revvity). CDK/cyclin enzymes were preincubated with test compounds for 20 min in 5 µL assay buffer (50 mM HEPES, 1 mM EGTA, 10 mM MgCl_2_, 0.01% Tween 20, 2mM DTT). Reactions were initiated by addition of 5 µL of 2x ATP and 2x ULight™-eIF4E-binding protein 1 (Thr37/46) peptide substrate (Revvity, TRF0128-M) and incubated for 60 min at room temperature. Reactions were terminated with 5 µL 4x EDTA, followed by addition of 5 µL of 4 x LANCE Ultra Europium-anti-phospho-eIF4E-binding protein 1 (Thr37/46) antibody (Revvity TRF0216-M) and incubated for 60 min at room temperature. The final concentrations of reagents in the activity assay were 1 mM ATP, 50 nM ULight™-eIF4E-binding protein 1 (Thr37/46) Peptide substrate, 1 % DMSO, 10 mM EDTA, and 0.5 nM LANCE Ultra Europium-anti-phospho-eIF4E-binding protein 1 (Thr37/46), and either 1 nM CDK4/CCND1, 0.25 nM CDK6/CCND3, 2 nM CDK2/CCNE1, or 0.5 nM CDK1/CCNB1.

TR-FRET signals were measured on an Envision plate reader (Perkin Elmer) with excitation at 340 nm and emission at 615 and 665 nm. The TR-FRET signal was calculated as the background subtracted 665/615 nm emission ratio. Percent inhibition was calculated relative to DMSO controls, and IC50 values were determined by fitting the concentration-response data to a four-parameter logistic nonlinear regression model.

### Cell lines and culture

T47D (RRID:CVCL_0553), ZR751 (RRID:CVCL_0588), REH (RRID:CVCL_1650), and HCC1428 (RRID:CVCL_1252) human breast cancer cell lines were obtained from ATCC and maintained in RPMI-1640 media supplemented with 10% fetal bovine serum (FBS) at 37°C in 5% CO_2_. EFM19 cells (RRID:CVCL_0253) were obtained from DSMZ and maintained in RPMI-1640 media supplemented with 10% FBS at 37°C in 5% CO_2_. MCF7 cells (RRID:CVCL_0031) were obtained from ATCC and maintained in DMEM media supplemented with 10% FBS and insulin. CAMA1 cells (RRID:CVCL_1115) were obtained from ATCC and maintained in EMEM media supplemented with 10% FBS. MV411 cells (RRID:CVCL_0064) were obtained from ATCC and maintained in IMDM media supplemented with 10% FBS. Isogenic RB1-deleted (RB-del) cell lines were generated by CRISPR-Cas9 genome editing using sgRNAs targeting exon 3 of *RB1* as published in our previous studies [23, 24]. The deletion was confirmed by immunoblot analysis. CDK6-overexpressing (CDK6 OE) T47D cells were generated by lentiviral overexpression vector, pLX304 containing the ORF clones for *CDK6*. The transduced cells were selected in the presence of blasticidin and CDK6 expression was confirmed by western blotting. All lines were routinely tested for mycoplasma contamination.

### CyQUANT cell proliferation assay

For end-point cell proliferation assays, cells were seeded in 384-well plates (Corning, 3571) in 40uL of cell culture medium at the following seeding densities: T47D (1,000 cells/well), MCF7 (600 cells/well), CAMA1 (1,200 cells/well), ZR751 (900 cells/well), EFM19 (1,400 cells/well), HCC1428 (1,800 cells/well), REH (1,500 cells/well), and MV411 (1,200 cells/well). CyQUANT Direct Proliferation Assay was performed according to manufacturer’s instructions (Invitrogen, C35012) after 6 days of compound treatment. Percent inhibition was determined relative to DMSO to determine IC50 values.

### Phospho-RB Ser780 AlphaLISA assay

T47D cells (6,000 cells/well) or REH cells (12,500 cells/well) were seeded in a 384-well plate in the presence of compound. RB phosphorylation was determined using the AlphaLISA SureFire Ultra Phospho-Rb (Ser780) Detection Kit (Revvity, ALSU-PRB-C500) after 24h of compound treatment according to manufacturer’s instructions.

### Live-cell Incucyte proliferation assays

For live-cell proliferation assays, cells stably expressing H2B-GFP were seeded in 96-well plates at 3,000–5,000 cells per well in 200 µL medium. After 24 hours, compounds were added at the indicated concentrations in technical triplicate. Plates were imaged every 4 hours using an Incucyte S3 Live-Cell Analysis System (Sartorius, RRID:SCR_023147), and proliferation was quantified as normalized GFP count. For colony formation assays, cells were seeded at low density (500 per well in 6-well plates) in the presence of vehicle (DMSO) or ALK-13743 (250 nM) and maintained for 34 days with medium and compound refreshed every 3 days; colonies were fixed and stained with crystal violet.

### Spheroid growth assay

To examine the impact of CDK4 inhibition on spheroid growth, T47D cells were seeded in 50% Matrigel/well in a 96-well TC dish (1500 cells/well) as described in our previous studies [24, 25]. The cells were allowed to form spheroids for 24 hours and exposed to DMSO and ALK-13743. The growth of spheroids was monitored in real-time using the Incucyte S3 system, based on the change in object area.

### Phalloidin staining

MCF7-WT and MCF7-RB-del cells were seeded onto coverslips and treated with ALK-13743 for 5 days. Following treatment, cells were fixed with 3.7% formaldehyde for 10 minutes and permeabilized using buffer containing 5% BSA and 0.5% Triton X-100. Cells were subsequently blocked with immunofluorescence buffer containing 5% BSA and 0.2% NP-40 for 5 minutes, followed by incubation with Alexa Fluor 568-conjugated phalloidin for 1 hour at room temperature. Nuclei were counterstained with DAPI, and fluorescence images were acquired using an EVOS microscope.

### HSPC myeloid maturation

Primary human CD34+ hematopoietic stem and progenitor cells (HSPCs) isolated from bone marrow (STEMCELL Technologies, 70002) were thawed and allowed to recover overnight at 37°C in 5% CO_2_ in IMDM supplemented with 10% FBS and StemSpan CC100 (STEMCELL Technologies, 2690). The next day, cells were treated with compound and differentiated with StemSpan CC100, 10 ng/mL GMCSF (STEMCELL Technologies, 78140) and 10 ng/mL G-CSF (STEMCELL Technologies, 78138). Media and compounds were refreshed every 2-3 days to maintain optimal cell density. Myeloid maturation was evaluated after 11 days of differentiation by flow cytometry staining and quantification of CD11b (Sigma, FCMAB178P, RRID:AB_10806342), CD45 (Invitrogen, 48-0459-42, RRID:AB_2016677), CD13 (Invitrogen, 11-0138-42, RRID:AB_11043278), and CD34 (Invitrogen, 17-0349-42, RRID:AB_2016672) surface markers. The percentage of CD11b+ cells was determined for each condition and normalized relative to that of DMSO samples to calculate IC50.

### Drug combination screening

A library of 307 agents (at 50 nM each, covering AKT, CDK, EGFR, ER, MEK, mTOR, PI3K, and other target classes) was screened in combination with ALK-13743 in T47D-WT and T47D-RB-del cells using the Incucyte proliferation platform. Confirmatory dose-response matrix experiments for ALK-13743 plus giredestrant or INX-315 were performed across 4×4 concentration matrices in MCF7 cells.

### Immunoblotting

Cells were treated for 24 hours, lysed in RIPA buffer supplemented with protease and phosphatase inhibitors, and resolved by SDS-PAGE on 4–12% Bis-Tris gels as described in our previous studies [26]. After transfer to PVDF membranes, blots were probed with antibodies against pRB (S807/811, #8516, Cell Signaling Technology, RRID:AB_11178658), CDK6 (#3136S, CST, RRID:AB_2229289), Cyclin A (#4656, CST, RRID:AB_2071958), Cyclin B1 (#12231, CST, RRID:AB_2783553), Cyclin D1 (#2978, CST, RRID:AB_2259616), β-Actin (#3700, CST, RRID:AB_2242334), and GAPDH (#5174, CST, RRID:AB_10622025 and #60004-1-Ig, Proteintech, RRID:AB_2107436). HRP-conjugated secondary antibodies and ECL detection were used for visualization.

### Xenografts

Six-to eight-week-old BALB/c nude mice were implanted subcutaneously at the left flank with estrogen pellets (0.36 mg / 60 days) 3 days prior to cell inoculation. T47D cells (Wuxi) were subcutaneously injected into the right flank at a density of 10×10^6^ cells per mouse. Once tumors reached ∼165 mm^3^, mice were randomized to treatment twice daily by oral gavage with vehicle (0.5% w/v methylcellulose A4M and 1% Tween 80 in water), 60 mg/kg atirmociclib, or 60 mg/kg ALK-13743. Tumors were measured twice a week with digital calipers. Mice were harvested either 3 hours or 12 hours following the last treatment to assess drug pharmacokinetics and pharmacodynamics. Sex as a biological variable was not assessed in the current study as only female mice were used due to the prevalence of females being impacted by breast cancer. Euthanasia was conducted by carbon dioxide inhalation followed by cervical dislocation.

### Ethics and approval

Animal care, treatment, and sacrifice were approved under protocol number 1384 M by the Institutional Animal Care and Use Committee (IACUC) at Roswell Park in accordance with the NIH Guide for the Care and Use of Laboratory Animals.

### Statistical analysis

Statistical analyses were performed in GraphPad Prism 10 (RRID:SCR_002798). Differences in growth rates between treatment conditions were evaluated by one-way ANOVA with Dunnett post-hoc testing. Significance thresholds: *p < 0.05, **p < 0.01, ***p < 0.001. Bliss synergy scores were reported as the mean of biological triplicates. All proliferation experiments were performed in at least three independent biological replicates. All *in vivo* analyses were conducted using three biological replicates. Power analyses were conducted prior to experimentation to identify the minimum number of mice required to sufficiently identify biological differences between treatment cohorts. Investigators were not blinded to treatment cohorts.

### Data availability

The data presented in this study are available from the corresponding author upon request.

## Results

### ALK-13743 is a potent and selective CDK4 inhibitor

ALK-13743 was developed through structure-guided medicinal chemistry as is described in detail in its patent filing[22] (Figure 1A). Biochemical Time-Resolved Fluorescence Resonance Energy Transfer (TR-FRET) profiling using 1 mM ATP confirmed that ALK-13743 functions as a potent and selective CDK4 inhibitor; ALK-13743 displayed an IC50 of 33 nM for CDK4/Cyclin D1, which was 3 times more selective than its IC50 for CDK6/Cyclin D3 (99 nM), 56 times more selective than its IC50 for CDK2/Cyclin E1 (1854 nM), and over 300 times more selective than its IC50 for CDK1/Cyclin B1 (>10000 nM) (Figure 1B). In cellular proliferation analyses, ALK-13743 displayed nanomolar potency comparable to atirmociclib (PF-07220060) in CDK4-dependent HR+ breast cancer cell lines (T47D, MCF7, CAMA1, ZR751, EFM19, HCC1428), while showing limited inhibition of CDK6-dependent cell lines (REH, MV411, HSPC) (Figure 1C). Dose response curves from T47D (blue) and REH cells (red) for inhibition of phosphorylated RB (left) and cell proliferation (right) revealed T47D selective inhibition when treated with atirmociclib or ALK-13743, while palbociclib had similar effects in both cell lines (Figure 1D); quantification of phosphorylated RB at serine residue 870 IC50 from Figure 1D is shown in Figure 1E. With CDK6 inhibitor-associated hematological toxicities being a central reason for the development of better tolerated CDK4-selective inhibitors, hematopoietic stem and progenitor cell (HSPC) myeloid maturation was further assessed across the three compounds. While palbociclib displayed comparable IC50 values in both T47D (78 nM) and HSPC cells (107 nM), ALK-13743 displayed robust selectivity for T47D cells (31 nM) and spared HSPC cells (831 nM) to a greater extent than CDK2/4/6 inhibitor RGT-419B (38 nM; 174 nM) and atirmociclib (50 nM; 629 nM) (Figure 1F).

**Figure 1.**
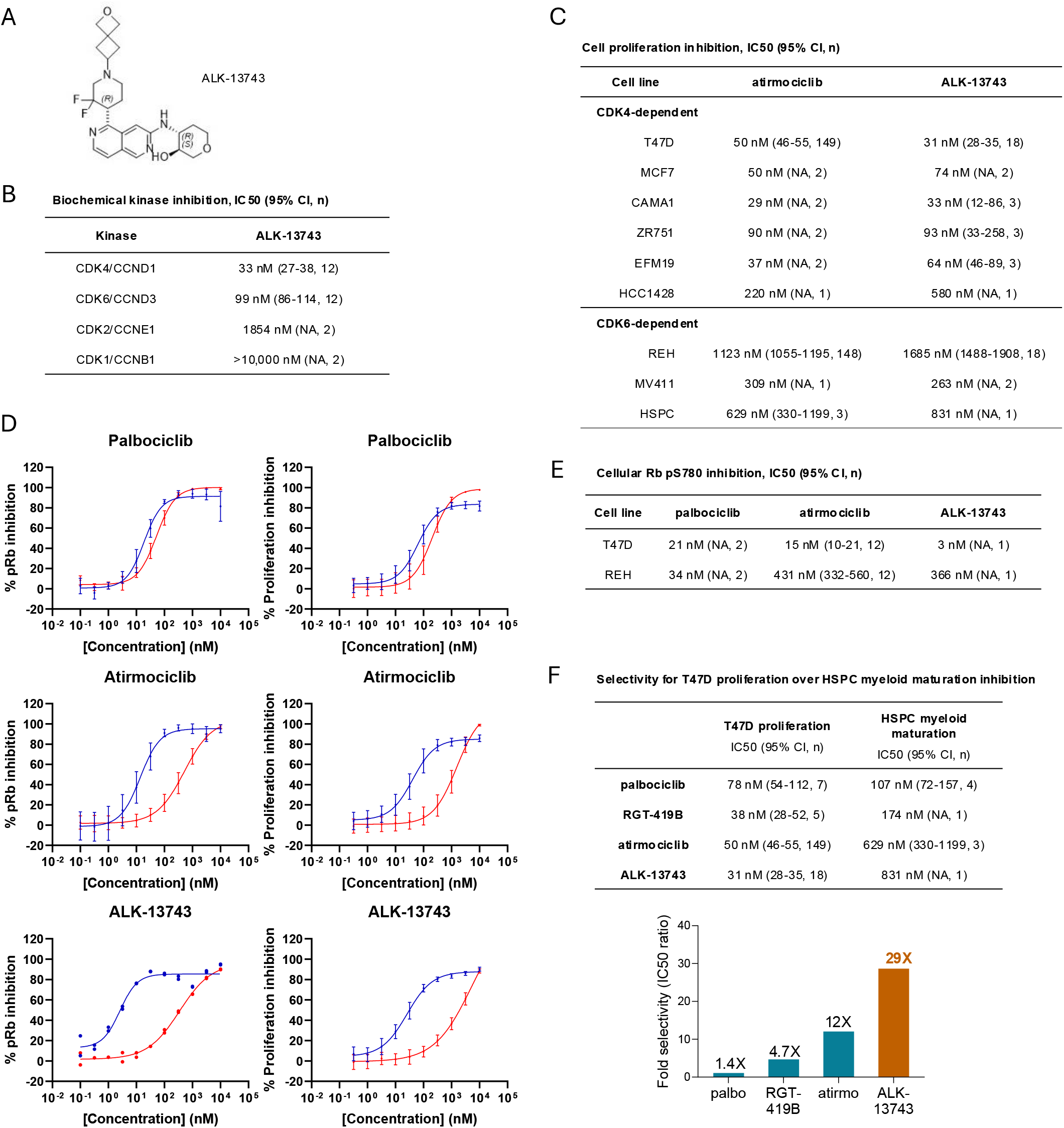
Chemical characterization and CDK4 selectivity of ALK-13743. (A) Chemical synthesis and structure of ALK-13743. (B) TR-FRET kinase selectivity profiling of ALK-13743 demonstrating selective inhibition of CDK4 over CDK6, CDK2, and CDK1. (C) Cell proliferation IC50 determination for ALK-13743 compared to the reference CDK4-selective inhibitor atirmociclib across various CDK4 vs CDK6-dependent cell lines. (D) Dose response curves of palbociclib, atirmociclib, and ALK-13743 in CDK4-dependent T47D (blue) and CDK6-dependent REH cells (red) based upon phosphorylated RB alphaLISA (left) and CyQUANT cell proliferation (right). (E) Combined dose response IC50 values for inhibition of phosphorylated serine residue 780 of RB with treatment of ALK-13743, palbociclib, or atirmociclib in the ER+ breast cancer cell line T47D compared to the hematopoietic REH cell line. (F) Quantified selectivity of palbociclib, RGT-419B, atirmociclib, and ALK-13743 for T47D proliferation and HSPC cell line myeloid maturation.

### Intracellular determinants of response to ALK-13743

To define the cellular response to intracellular CDK4 inhibition, we examined the efficacy of ALK-13743 in the CDK4-dependent ER+ breast cancer models MCF7 and T47D. ALK-13743 potently suppressed proliferation in both lines, with activity evident at concentrations as low as 125 nM (Figure 2A).

**Figure 2.**
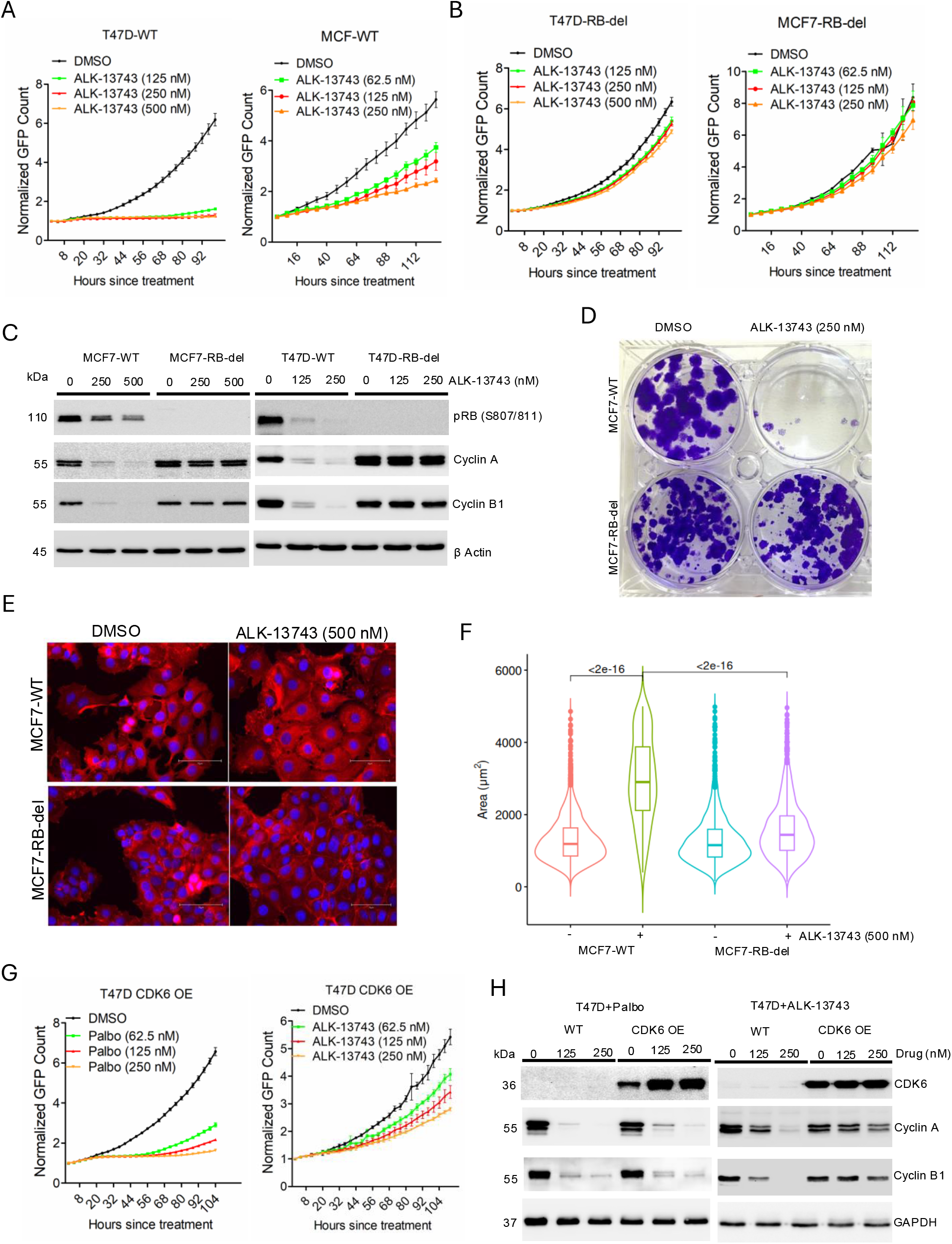
RB pathway dependence of ALK-13743 antiproliferative activity. (A) Incucyte live-cell proliferation curves of T47D-WT and MCF7-WT cells treated with ALK-13743 at the indicated concentrations. Error bars represent mean and SD from triplicates. Experiment was done at three independent times. (B) Effect of ALK-13743 on the proliferation of T47D-RB-del and MCF7-RB-del cells. Error bars were determined based on mean and SD. Experiment was done at 3 independent times. (C) Biochemical analysis on T47D-WT, MCF7-WT, T47D-RB-del and MCF7-RB-del cells following 48-hour treatment with ALK-13743 at the indicated concentrations. (D) Crystal violet colony formation assay of MCF7-WT and MCF7-RB-del cells treated with ALK-13743 versus DMSO control. Colonies were treated for 34 days. (E) Representative images from Phalloidin staining (red) on the MCF7-WT and MCF7-RB-del cells following 48-hour treatment with ALK-13743. (F) Violin plots indicating the cell size distribution from MCF7-WT and MCF7-RB-del cells following 5-day treatment with ALK-13743. P values were determined based on student t-test. (G) Differential effects of palbociclib and ALK-13743 on the proliferation of T47D-CDK6 OE cells. Error bars represent mean and SD from triplicates. Experiment was done at three independent times. (H) Immunoblot analysis on the indicated proteins from T47D-WT and T47D CDK6 OE cells treated with palbociclib or ALK-13743.

It has been shown that loss of RB, the preferential substrate of CDK4 and CDK6, renders cells resistant to both genetic depletion of CDK4/6 and pharmacological inhibition with palbociclib [24, 27, 28]. To determine whether this dependency also governs the response to ALK-13743, we used isogenic MCF7 and T47D derivatives harboring genetic deletion of RB (MCF7-RB-del and T47D-RB-del). The cytostatic effect of ALK-13743 was abrogated in both RB-deleted lines, indicating that its antiproliferative activity is strictly RB-dependent (Figure 2B). Consistent with this, biochemical analysis showed that ALK-13743 inhibited RB phosphorylation and suppressed the E2F-regulated cell cycle proteins cyclin A and cyclin B1 in parental MCF7 and T47D cells (Figure 2C). In the RB-deleted setting, cyclin A and cyclin B1 levels were maintained despite CDK4 inhibition, confirming that the cytostatic effect of ALK-13743 is mediated through RB activation (Figure 2C). Long-term colony formation assays yielded the same result, with RB loss conferring resistance to ALK-13743 (Figure 2D). The cytostatic response to ALK-13743 was accompanied by a marked change in cell morphology, with phalloidin staining revealing cell enlargement (Figure 2E & 2F). This effect was also RB-dependent as the MCF7-RB-del cells showed no increase in cell size, indicating that enlargement is a consequence of proliferative arrest rather than a distinct drug effect (Figure 2E & 2F).

To determine whether ALK-13743 preferentially targets CDK4 over CDK6 within cells, we ectopically overexpressed CDK6 in T47D cells [21]. CDK6 overexpression in T47D cells (T47D CDK6 OE) had limited effect on sensitivity to palbociclib, which inhibits both CDK4 and CDK6, but largely abrogated the antiproliferative effect of ALK-13743 (Figure 2G). Biochemical analysis further supported that following CDK6 overexpression, palbociclib retained the ability to suppress cyclin A and cyclin B1 to an extent comparable with that in parental T47D cells, whereas ALK-13743 had only a modest effect on these cell cycle regulatory proteins in T47D CDK6 OE cells (Figure 2H). These data indicate that the compensatory activity conferred by CDK6 overexpression can be limited by palbociclib but not by ALK-13743, establishing CDK4 as the preferential cellular target of ALK-13743. Together, these findings demonstrate that the antiproliferative activity of ALK-13743 is on-target and RB-dependent, and that it is selectively bypassed by CDK6.

### ALK-13743 exhibits broad combination activity in RB-proficient cells

Given the established clinical rationale for combining CDK4/6 inhibitors with endocrine therapy, we assessed the efficacy of ALK-13743 in combination with giredestrant, a recently developed selective estrogen receptor degrader (SERD)[15, 29, 30]. The combination produced a synergistic antiproliferative interaction in both MCF7 and T47D cells (Figure 3A), and the single-agent activity of ALK-13743 was markedly enhanced by giredestrant, yielding growth arrest (Figure 3B). This enhanced cytostatic effect was accompanied by cooperative inhibition of RB phosphorylation and downregulation of the cell cycle proteins cyclin A and cyclin B1 in both lines, relative to either single agent (Figure 3C). To determine whether this combination effect requires RB, we compared the response of parental and isogenic MCF7-RB-del cells. In contrast to parental cells, cyclin A and cyclin B1 levels were maintained across all treatment conditions in MCF7-RB-del cells, whereas the parental cells showed cooperative suppression (Figure 3D).

**Figure 3.**
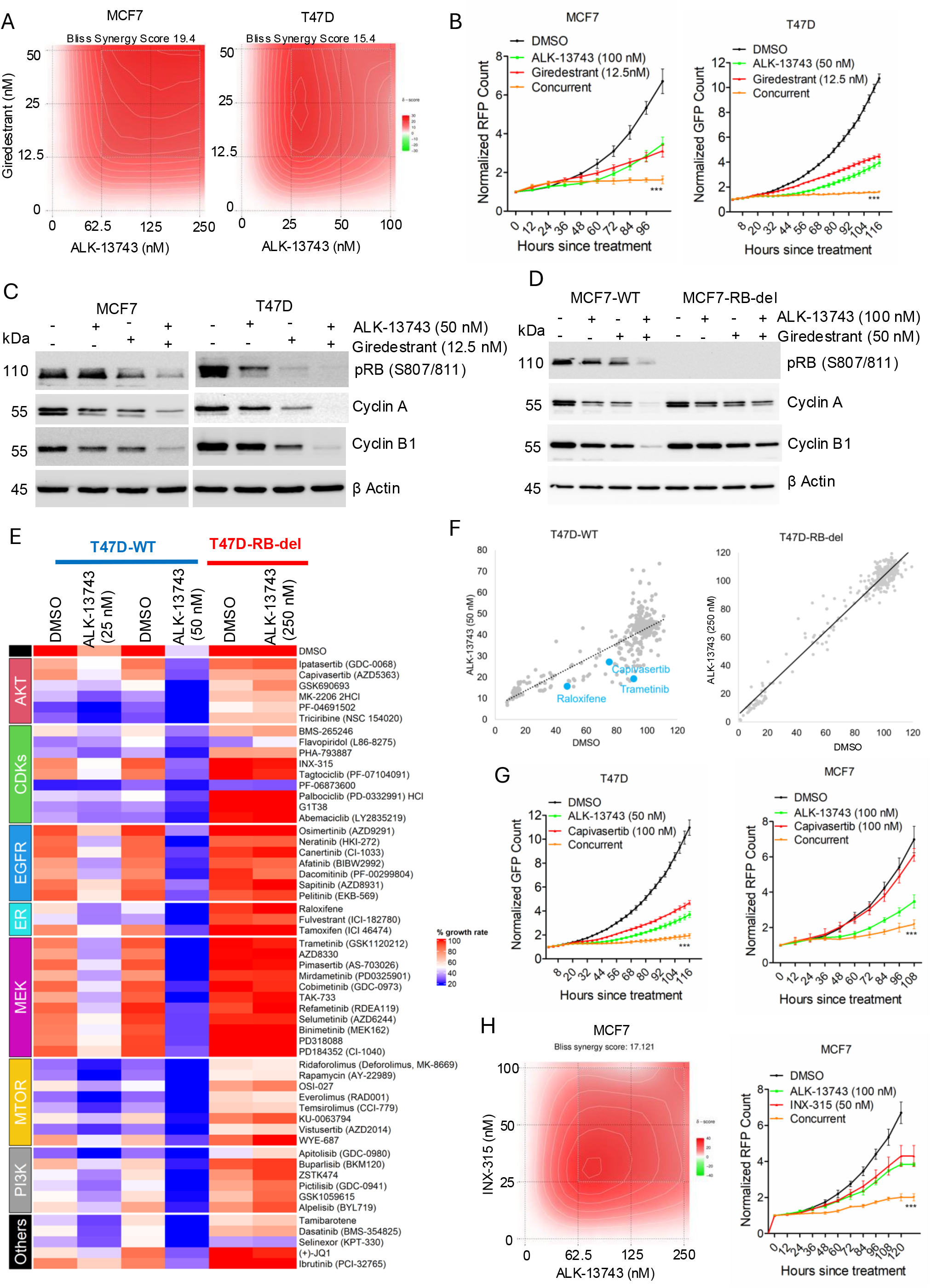
Broad RB-dependent combination activity of ALK-13743 with endocrine and PI3K-pathway inhibitors. (A) Synergistic interaction between giredestrant and ALK-13743 in MCF7 and T47D cells. The Bliss synergy score was calculated using Synergy Finder. (B) Effect of giredestrant in combination with ALK-13743 on the proliferation of MCF7 and T47D cells. Error bars represent mean and SD from triplicates. Experiment was done at three independent times. *** p<0.0001 as determined by 2-way ANOVA. (C) Biochemical analysis in MCF7 and T47D cells following the treatment with giredestrant in combination with ALK-13743 for 48 hours. (D) Differential effect of giredestrant in combination with ALK-13743 on the cell cycle proteins between MCF7-WT and MCF7-RB-del cells. (E) Heat map illustrating the combined effect of ALK-13743 with the indicated drugs in T47D-WT and T47D-RB-del cells based on cell proliferation. (F) Scatter plot of T47D-WT and RB-del growth rate for ALK-13743 treatment in combination with DMSO versus the drug library. (G) Incucyte live-cell proliferation curves for T47D and MCF7 cells following the treatment with ALK-13743 in combination with capivasertib. (H) Bliss synergy plot for the combined treatment of MCF7 cells with ALK-13743 and INX-315 at the indicated concentrations. Effect of ALK-13743 in combination with INX-315 on the proliferation of MCF7 cells. Error bars represent mean and SD from triplicates. Experiment was done at three independent times. *** p<0.0001 as determined by 2-way ANOVA.

To define the broader combinatorial landscape of ALK-13743 and the role of RB within it, we performed a drug screen using a 307-compound library spanning AKT, CDK, EGFR, ER, MEK, mTOR, PI3K, and additional target classes, scoring each combination by effect on proliferation rate (Figure 3E). In T47D cells, ALK-13743 combined with multiple classes of targeted agents — including endocrine therapies and PI3K, mTOR, and AKT inhibitors — and produced cooperative growth arrest, while in T47D-RB-del cells, this combinatorial benefit was uniformly lost (Figure 3E). Consistent with these findings, correlation analysis comparing drug responses in the absence (DMSO) or presence of ALK-13743 revealed a broad enhancement of drug efficacy following CDK4 inhibition in T47D-WT cells. However, this effect was markedly diminished in T47D-RB-del cells, indicating that RB loss compromises the ability of CDK4 inhibition to enhance responses to other targeted agents (Figure 3F). This indicates that functional RB is required not only for single-agent CDK4 inhibitor efficacy but also for therapeutic benefit from these combinations.

We validated these screen results with representative agents. Combination of ALK-13743 with the AKT inhibitor capivasertib significantly enhanced antiproliferative activity in both MCF7 and T47D cells (Figure 3G). The screen also identified cooperative activity with CDK2 inhibitors, consistent with CDK2 acting downstream of CDK4. Validation with the selective CDK2 inhibitor INX-315 confirmed a synergistic interaction with ALK-13743 and produced durable proliferative arrest (Figure 3H). Overall, the data demonstrates that therapeutic efficacy of ALK-13743 could be enhanced through different combinatorial approaches involving targeted therapeutics in an RB proficient setting.

### Functional role of CDK6 on impacting the therapeutic benefits of ALK-13743

To characterize the activity profile of ALK-13743 in the context of CDK6-mediated resistance, we compared combination responses in parental and CDK6-overexpressing T47D cells. ALK-13743 combined with giredestrant produced an enhanced antiproliferative effect in parental T47D cells; however, this response was attenuated in T47D-CDK6-OE cells (Figure 4A). Biochemical analysis supported this result: in CDK6-overexpressing cells, RB phosphorylation and expression of E2F-regulated cell cycle proteins were maintained despite combined treatment with ALK-13743 and giredestrant (Figure 4B).

**Figure 4.**
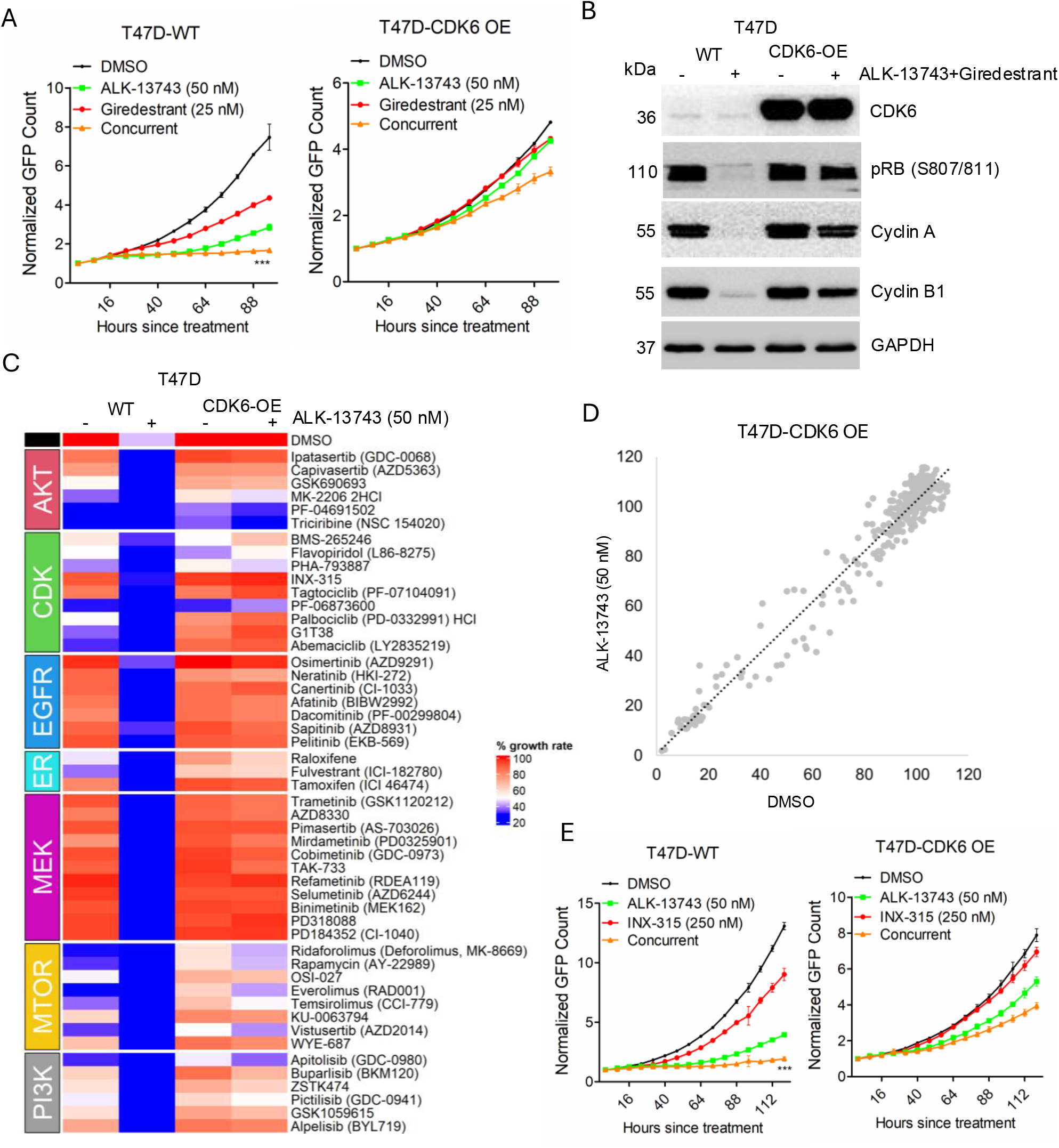
CDK6 overexpression abrogates T47D growth arrest when treated with ALK-13743 alone and in combination with drug library. (A) Effect of ALK-13743 in combination with giredestrant on the proliferation of T47D-WT and T47D-CDK6 OE cells. Error bars represent mean and SD from triplicates. Experiment was done at three independent times. (B) Western blot analysis on T47D-WT and T47D-CDK6 overexpressing (OE) cells treated with ALK-13743 (50 nM) in combination with giredestrant (25 nM) for 48 hours. (C) Heatmap depicting percent growth rate of T47D-WT and T47D CDK6 OE cells treated with DMSO, and ALK-13743 in combination with a 307-drug library. (D) Scatter plot of T47D CDK6 OE growth rate for ALK-13743 treatment in combination with DMSO versus the drug library. (E) Effect of ALK-13743 in combination with INX-315 on the proliferation of T47D-WT and T47D-CDK6 OE cells. Error bars represent mean and SD from triplicates. Experiment was done at three independent times. *** p<0.0001 as determined by 2-way ANOVA

We next performed a combinatorial drug screen in T47D-CDK6-OE cells alongside parental T47D comparators, using ALK-13743 at 50 nM. While ALK-13743 produced potent combinatorial growth arrest in parental cells, CDK6 overexpression broadly attenuated the combination benefit across all drug classes tested (Figure 4C, 4D). We validated this result with the CDK2 inhibitor INX-315: co-targeting CDK4 and CDK2 cooperatively inhibited proliferation in parental T47D cells, whereas CDK6 overexpression mitigated the response to the combination (Figure 4E), consistent with the screen.

### ALK-13743 displays strong anti-tumor activity in breast cancer xenograft models

ALK-13743 activity was then assessed in 3D culture and in preclinical mouse models. ALK-13743 drastically reduced T47D spheroid growth (Figure 5A). Activity was then assessed alongside atirmociclib *in vivo* in NSG mice implanted with orthotopic T47D tumors. While tumor growth inhibition was observed in mice treated with atirmociclib, mice treated with ALK-13743 displayed tumor regression, with no observed adverse effects as measured by body weight (Figure 5B). As was observed *in vitro*, ALK-13743 treatment resulted in substantial reduction of pRB, while pRB remained at appreciable levels in atirmociclib-treated tumors (Figure 5C). Additionally, pRB remained low 12 hours post ALK-13743 treatment, whereas pRB levels slightly rebounded at 12 hours post atirmociclib treatment. These findings are consistent with the *in vitro* studies and provide proof-of-concept for the therapeutic potential of ALK-13743 in ER+, RB proficient breast cancer.

**Figure 5.**
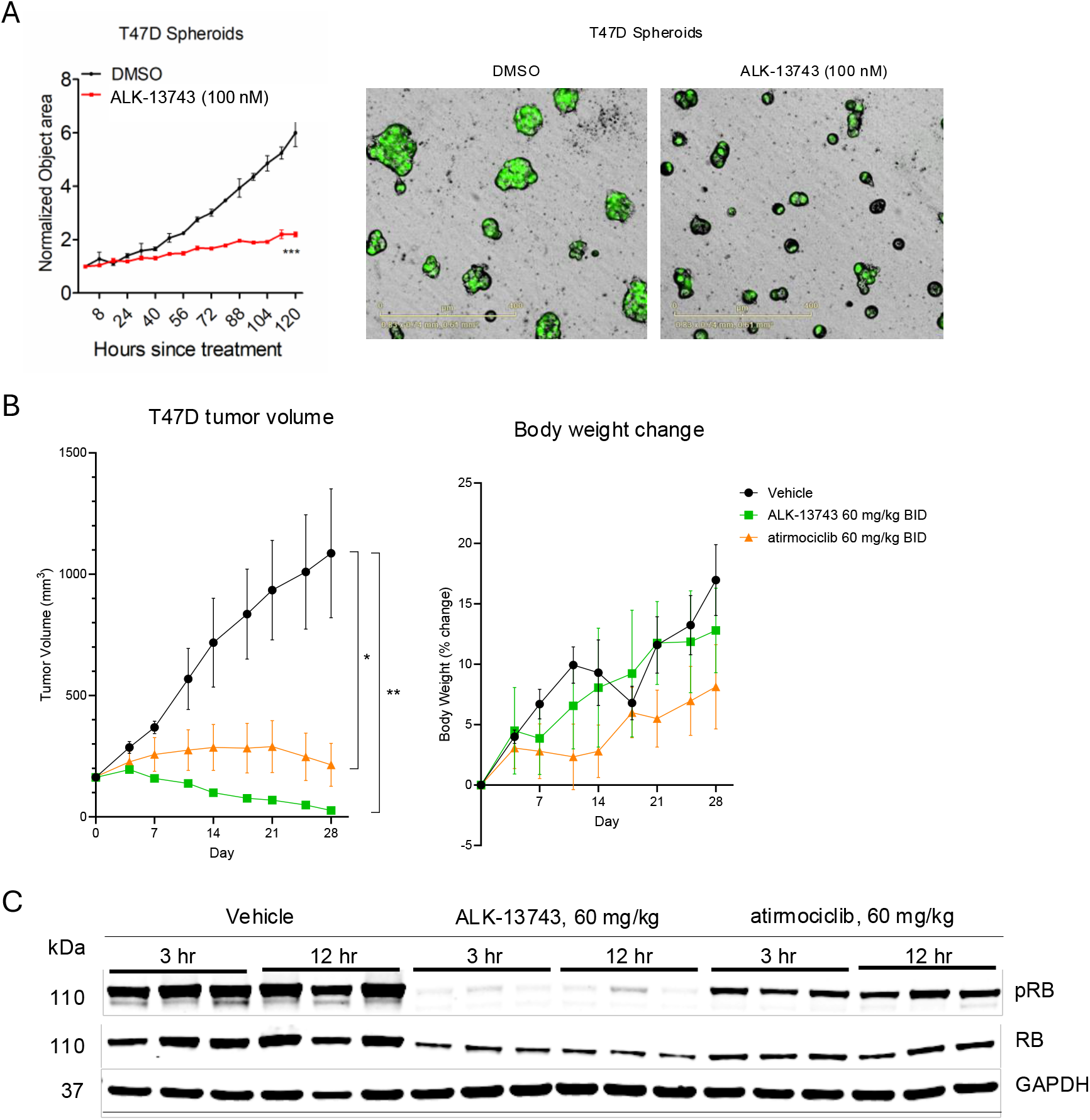
3D culture and *in vivo* efficacy of ALK-13743 in HR+ breast cancer models. (A) T47D spheroid growth when treated with DMSO or ALK-13743, with associated image stills. (B) *In vivo* tumor volume and body weight change of xxT47D xenograft-bearing BALB/c nude mice treated with vehicle, ALK-13743, or atirmociclib 60 mg/kg twice daily. ** p<0.01 and * p<0.05 as determined by one-way ANOVA (C) Immunoblot analysis of phospho-RB, total RB, and GAPDH from excised xenografts at 3 hours or 12 hours following the final treatment with vehicle, atirmociclib, or ALK-13743.

## Discussion

Despite the success of CDK4/6 inhibitors in HR+ breast cancer, their toxicity profile has limited their use and likely efficacy in different disease contexts. Palbociclib and ribociclib produce grade 3/4 neutropenia in over 50% of patients[15, 31]. The observation that CDK4-selective inhibitors can spare hematopoietic cells while retaining antitumor activity was first suggested in preclinical models using genetic approaches and subsequently reinforced by preclinical studies of atirmociclib[18, 19]. In early-stage clinical results, atirmociclib produced substantially less neutropenia compared to dual CDK4/6 inhibitors [20, 21]. CDK4 and CDK6 are highly conserved with >70% amino acid identity and structural similarity. For selectivity, a series of molecules were developed to interact more potently with CDK4/Cyclin D1 vs. CDK6/Cyclin D3 complexes. This selectivity, which was validated in biochemical assays, was more pronounced in cell-based proliferation assays. Similarly, ALK-13743 exhibited a selectivity ratio superior to atirmociclib when assessing impact on HR+/HER2-breast cancer vs. hematopoietic stem cell myeloid maturation. These results suggest that the *in vivo* CDK4 and CDK6 holoenzyme may behave differently than the dimers employed for *in vitro* kinase assays.

Consistent with previous studies demonstrating the pharmacology of CDK4/6 inhibitors (e.g. palbociclib), ALK-13743 potently suppressed the proliferation of HR+/HER2-breast cancer cells, which was associated with the dephosphorylation of RB. While RB loss is not the sole driver of resistance to CDK4/6 inhibitors clinically, it is observed in ∼10% of tumors following clinical progression in the context of randomized clinical trials. The strict RB dependence of ALK-13743 activity is consistent with prior characterizations of CDK4/6 inhibitors and underscores RB loss as a universal resistance mechanism to drugs targeting this axis [7, 10, 23]. In addition to RB loss, CDK6 is implicated in driving therapeutic resistance both through gene amplification and via deregulated expression. The present data, using ectopic CDK6 overexpression, indicate that elevated CDK6 compensates for selective CDK4 inhibition far more effectively than for dual CDK4/6 inhibition.

A key feature of CDK4/6 inhibitors is their ability to be combined effectively with other targeted therapies. As shown here, ALK-13743 broadly cooperates with agents similar to that observed with the CDK4/6 inhibitor palbociclib and CDK4 inhibitor atirmociclib. This cooperation was confirmed with endocrine therapy and AKT inhibitors that are FDA-approved for the treatment of HR+/HER2-breast cancer[30, 32]. Since CDK4 inhibitors are currently being tested clinically post-progression on CDK4/6 inhibitors, we evaluated the influence of RB loss and CDK6 overexpression on combinatorial activity with palbociclib, atirmociclib, and ALK-13743. Both CDK6 overexpression and loss of RB yielded resistance to all CDK4/6 targeting drugs as expected. However, no combination was effective at cooperating with the CDK4/6 or CDK4 inhibitors in the context of this form of therapeutic resistance. These findings suggest that knowledge of the drivers of resistance could help to inform utilization of select combinations moving forward.

In summary, ALK-13743 represents a structurally novel, potent, and highly CDK4-selective inhibitor with a mechanistically consistent antiproliferative profile in HR+ breast cancer models. Its combination activity with endocrine and PI3K-pathway agents establish a strong preclinical rationale for clinical investigation. ALK-13743 joins a small but growing group of CDK4-selective inhibitors—alongside atirmociclib and other emerging agents—that may define a new treatment paradigm of highly selective cell cycle targeting with reduced myelosuppression and improved combinability in HR+/HER2-breast cancer and other malignancies.

## Acknowledgements

The authors thank all members of the laboratory group and colleagues in the discussion and preparation of the manuscript. We thank Aleksia Therapeutics for providing ALK-13743. We thank the OnKure Therapeutics team for their assistance and oversight carrying out the *in vivo* experiments. We thank the Aleksia Therapeutics medicinal chemistry team of Claude Quesnelle, Rachelle Quach, and Sara Coulup for their work in designing and developing ALK-13743. Claude generative AI was used in the writing process to review and revise the manuscript text for completeness and clarity.

## Funding

This work was supported by grants from the National Institutes of Health, National Cancer Institute (CA267467 and CA211878) and was also supported by Roswell Park Comprehensive Cancer Center and National Cancer Institute grant P30CA016056. The content is solely the responsibility of the authors and does not necessarily represent the official view of the National Institutes of Health.

## Author contributions

Study concept and design: VK, JS, RH, AB, JB, AH, AP, EP, ESK and AKW. Acquisition of data: VK, JS, RH, AB, JB, KL and JW.

Analysis and interpretation of data: VK, TNO, JS, RH, AB, AH, JB, AP, KL, EP, ESK and AKW. Study supervision: VK, EP, ESK and AKW.

## Competing interests

Aleksia Therapeutics provided ALK-13743 to the Roswell Park investigators for this study. JS, RH, AB, JB, AH, AP, and EP are employees of Aleksia Therapeutics. ESK has sponsored research funded by Aleksia Therapeutics, Blueprint Medicines and Bristol Myers Squibb and is a member of the Cancer Cell Cycle-LLC consulting enterprise. AKW has sponsored research funded by Aleksia Therapeutics, Blueprint Medicines and Bristol Myers Squibb. All other authors declare no potential conflicts of interest.

## Notes

https://patentscope.wipo.int/search/en/detail.jsf?docId=WO2025240517&_cid=P22-MWDYEA-20599-1

### Competing Interest Statement

Aleksia Therapeutics provided ALK-13743 to the Roswell Park investigators for this study. JS, RH, AB, JB, AH, AP, KL, and EP are employees of Aleksia Therapeutics. ESK has sponsored research funded by Aleksia Therapeutics, Blueprint Medicines and Bristol Myers Squibb and is a member of the Cancer Cell Cycle-LLC consulting enterprise. AKW has sponsored research funded by Aleksia Therapeutics, Blueprint Medicines and Bristol Myers Squibb. All other authors declare no potential conflicts of interest.

### Summary of Updates

Primarily added an author that made substantial contributions to the design and development of ALK-13743 and acknowledged the rest of the medicinal chemistry team that assisted with drug development. Additionally corrected a misspelled author's name and made minor aesthetic changes to the figures.

